# The ESCRT-0 protein HRS regulates hepatocellular lipid droplet catabolism

**DOI:** 10.1101/2025.11.13.686840

**Authors:** Mathilda M. Willoughby, Ankit Shroff, Bridget E. Crossman, Micah B. Schott

## Abstract

Lipid droplets (LDs) are dynamic organelles that regulate lipid storage and metabolism pathways central to metabolic liver disease. LD turnover occurs in part through lysosomal catabolism (i.e. lipophagy) whereby LDs are thought to follow two distinct trafficking pathways: autophagosome-dependent *macro*lipophagy and the autophagosome-independent *micro*lipophagy. However, the molecular machinery that regulates these two distinct pathways, especially that of microlipophagy in mammalian cells, is poorly understood. In yeast, microlipophagy has been shown to rely on a protein family known as the endosomal sorting complex required for transport (ESCRT). Here, we used an ESCRT-specific RNAi library in hepatocytes which identified the ESCRT-0 protein hepatocyte growth factor receptor substrate (HRS) as a critical regulator of LD homeostasis. HRS depletion leads to significant LD accumulation which is not due to increased LD formation but from impaired LD catabolism. HRS-deficient cells retain lipolysis activity; however, they exhibit decreased LD targeting via microlipophagy, accompanied by compensatory increases in autophagosome targeting to LDs. In agreement with these findings, HRS knockdown suppressed mTOR signaling, boosted autophagosome formation, and reduced the degradation of autophagic cargo. Despite maintaining lysosome numbers, HRS knockdown raised lysosomal pH causing decreased autophagic degradative capacity and contributing to LD accumulation. Overall, these findings identify HRS as a modulator of LD turnover in mammalian cells, regulating lipophagy through lysosomal function.

**Significance Statement:**

- The regulatory molecular mechanisms of lipophagy are not clearly defined. This study identifies novel ESCRT proteins as regulators of LD homeostasis in several cell lines.
- In hepatocytes, we identified HRS specifically regulates LD catabolism, whereby HRS-dependent regulation of LDs is dual-faceted, affecting LD-lysosomal targeting and lysosomal function.
- Our findings are significant because they provide mechanistic insights into the role of ESCRT proteins in LD metabolism. Elucidating ESCRT-mediated lipophagy can potentially aid in developing novel targets to prevent aberrant lipid trafficking and utilization, particularly in the liver where LDs can accumulate and cause irreversible liver damage.

## Introduction

Lipid droplets (LDs) are highly dynamic cellular organelles composed of a neutral lipid core of triacylglycerols (TAGs) and sterol esters (SEs). This core is enclosed by a phospholipid monolayer embedded with a diverse and dynamic proteome. These proteins mediate essential functions, including maintaining the structural integrity of LDs and establishing contact sites with other organelles for lipid transfer, signaling, and metabolism. LD homeostasis, a central determinant of cellular energy status, is tightly regulated by the balance between biogenesis and catabolism. Disruption of this balance, leading to LD accumulation, contributes to multiple metabolic disorders, including metabolic dysfunction-associated steatohepatitis (MASH), alcohol-associated fatty liver disease (ALD), diabetes, and atherosclerosis.^1^ Thus, defining the molecular mechanisms regulating LD formation and turnover remains necessary.

LD biogenesis occurs at the endoplasmic reticulum (ER), where neutral lipids synthesized by enzymes such as diacylglycerol acyltransferase 1 (DGAT1) accumulate between the ER membrane leaflets and bud into the cytoplasm. Once released, LDs can grow and further enlarge through local TAG synthesis catalyzed by DGAT2. LD catabolism proceeds via two sequential pathways: cytosolic lipolysis and lysosome-mediated lipophagy.^2^ During lipolysis, cytoplasmic lipases, including adipose triglyceride lipase (ATGL), preferentially act on large LDs, liberating fatty acids (FAs) for β-oxidation and signaling. Lipophagy acts through trafficking the LDs to the lysosomes for degradation. Lipophagy was first identified as an autophagosome-dependent pathway (macrolipophagy), involving the sequestration of LDs into autophagosomes that subsequently fuse with lysosomes, where lysosomal acid lipase (LAL) degrades them.^3^ Following studies identified an autophagosome-independent form of lipophagy, termed microlipophagy, that was described in yeast and recently in Drosophila and mammalian cells, including hepatocytes,^4–6^ macrophages,^7^ and cardiomyocytes.^8^ Despite growing recognition, the molecular machinery that governs mammalian microlipophagy remains poorly defined.

Insights from yeast implicate the Endosomal Sorting Complexes Required for Transport (ESCRT) in microlipophagy regulation.^9–11^ ESCRT proteins are canonically known for directing endocytosed cargo into multivesicular bodies and lysosomes. They are organized into subcomplexes: ESCRT-0, –I, –II, –III, associated regulatory factors, and Bro1 domain–containing proteins.^12, 13^ Recent studies have found that ESCRT proteins VPS4A and VPS4B significantly affected LD accumulation in mouse hepatocytes^14^ while CHMP1 and CHMP4a negatively regulated LD catabolism in Drosophila.^15^

Here, we employed an RNAi screen of ∼70 known mammalian ESCRTs and ESCRT adaptors and measured total LD area/cell using high-content quantitative microscopy. These studies identified the ESCRT-0 protein hepatocyte growth factor receptor tyrosine kinase substrate (HGS, commonly called HRS) as a regulator of LD catabolism in hepatocytes. RNAi-mediated knockdown of HRS caused robust LD accumulation, not through enhanced LD biogenesis but due to impaired degradation. Mechanistic analyses revealed that HRS depletion did not significantly affect cytosolic lipolysis but reduced LD clearance through lipophagy. In HRS-depleted cells, we observed a profound accumulation of LDs within Lamp1-positive lysosomes. This was accompanied by a decrease in LD targeting by endosomal Rab5, a critical microlipophagy regulator, we well as a compensatory increase in LD targeting by LC3-positive autophagosomes and suppressed mTOR signaling. Despite increased autophagosome-LD targeting, HRS knockdown increased lysosomal pH, impairing their degradative capacity (i.e. decreased autophagy flux), ultimately leading to elevated LD levels. Together, these findings identify HRS as a critical mediator of LD turnover through coordinated regulation of lipophagy and lysosomal function, providing mechanistic insights into the role of ESCRT proteins in LD metabolism pathways.

## Results

### ESCRT proteins regulate lipid droplet levels in hepatocytes

To identify specific ESCRT proteins involved in LD homeostasis, we performed a targeted RNAi screen using Hep3B hepatoma cells. Following 72 h knockdown, cells were imaged using high-throughput epifluorescence microscopy. Total LD area/cell was quantified using CellProfiler image analysis. Lipolysis inhibition by adipose triglyceride lipase (ATGL) knockdown was used as a positive control for LD accumulation. Interestingly, the screen uncovered both positive and negative regulators of LD homeostasis (**Figure 1A and Table S1**). These proteins were not restricted to a single sub-family of ESCRTs but were from all four sub-groups (ESCRT-0, –I, –II, III) (**Figure S1A**). Among these hits, *Hrs* knockdown caused significant LD accumulation (**Figure 1B**) like that of ATGL knockdown. These findings were also consistent in HeLa cells (**Figure S1B, Table S2**).

**Figure 1.**
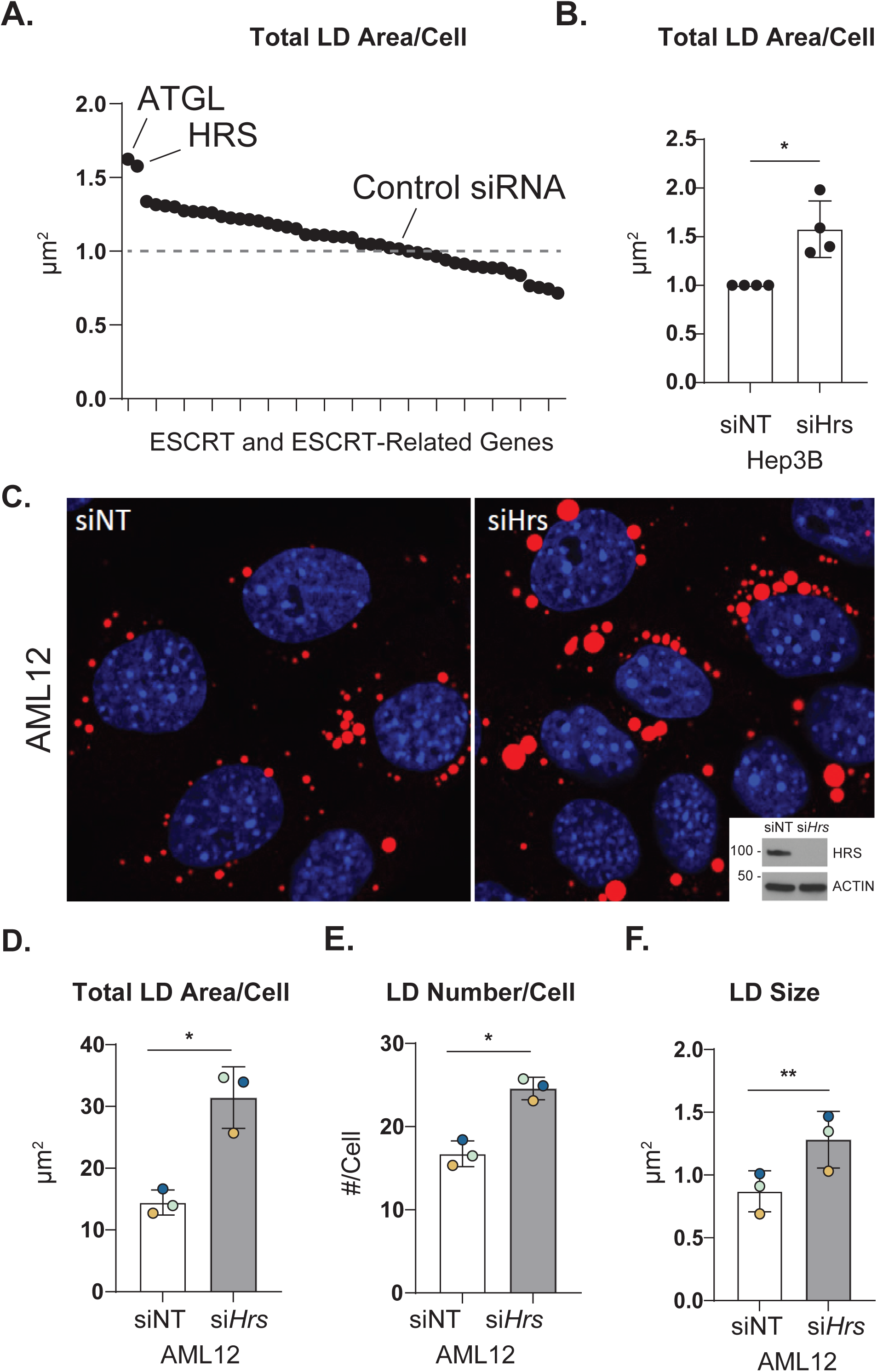
The ESCRT-0 protein HRS negatively regulates LD levels. **A**. Graph showing changes in total LD Area per cell from RNAi screening of ESCRT and ESCRT-related genes in Hep3B cells. **B.** Quantification of total LD area per cell in Hep3B cells upon *Hrs* knockdown. **C.** Confocal micrographs displaying ORO-stained LDs (red) and DAPI-stained nuclei (blue) in AML12 cells treated with control (siNT) or Hrs-targeting (si*Hrs*) siRNAs for 48 hours. Quantification of (**D**) total LD area per cell, (**E**) number of LDs per cell, and (**F**) LD size in cells from (**C**) using ImageJ. Data represent the mean ± SD from at least three independent trials, with each dot representing an individual trial. The statistical significance is indicated by asterisks using a two-tailed Student’s t-test for ratio-pairwise comparisons. \**P* < 0.05, \*\*\**P* < 0.001. Abbreviations: LD: Lipid Droplet; ESCRT: Endosomal Sorting Complex Required for Transport; ORO: Oil Red O.

### The ESCRT-0 protein HRS negatively regulates LDs

HRS is an ESCRT-0 protein known to bind phosphatidylinositol 3-phosphate (PI3P) at early endosome membranes via its FYVE domain. At the early endosome, it facilitates critical functions, including interacting with ubiquitinated cargo via its ubiquitin-interacting motif (UIM), sorting cargo for lysosomal degradation, and recruiting subsequent ESCRT sub-groups.^16–18^ A well-known characteristic of HRS-deficient cells is significantly enlarged endosomes (**Figure S1C**), suggesting HRS plays a role in endosome maturation.^19^ We observed an increase in LDs upon *Hrs* knockdown across multiple cell types. In addition to the initial findings from Hep3B cells, we found that LD area/cell was significantly increased following *Hrs* knockdown in AML12 normal mouse hepatocytes (**Figure 1C-F**), HeLa cells (**Figure S1D**), and HepG2 cells (**Figure S1E**).

### HRS-mediated LD accumulation is independent of LD biogenesis

LD accumulation is predominantly attributed to increased LD biogenesis and/or decreased LD degradation.^20^ To determine if HRS plays a role in LD biogenesis, we loaded control (siNT) and *Hrs* knockdown (si*Hrs*) AML12 cells with oleic acid (OA, 150 µM OA for 4 h) and simultaneously treated with DMSO vehicle control or inhibitors against DGAT1 and DGAT2 to prevent LD biogenesis. Our rationale was that if *Hrs* knockdown elevates LD levels by accelerating LD biogenesis, then HRS-deficient cells may exhibit a reduction in sensitivity to DGAT1/2 inhibitors (DGAT1/2i). However, both control and *Hrs* knockdown cells were equally responsive to DGAT1/2i, where simultaneous treatment of OA and DGAT1/2i abolished LD biogenesis in both siNT and si*Hrs* AML12 cells (**Figure 2A, B; Figure S2A, B**). This suggests that LD accumulation in HRS-deficient cells may occur due to decreased LD catabolism, rather than enhanced LD biogenesis.

**Figure 2.**
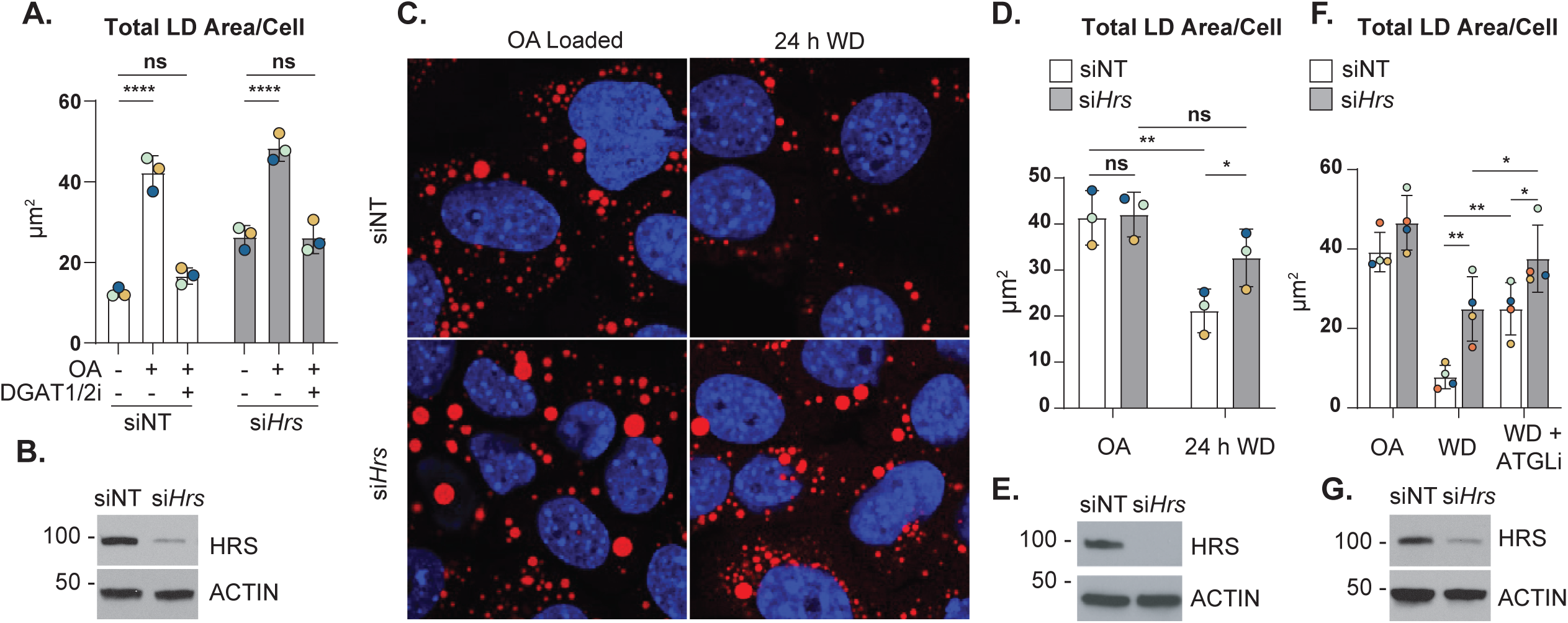
HRS-mediated LD accumulation is independent of LD biogenesis and lipolysis. **A.** Quantification of total LD area per cell in AML12 cells transfected with control (siNT) or *Hrs*-targeting (si*Hrs*) siRNAs and treated with OA, with or without DGAT1/2i. **B.** Immunoblot analysis confirming *Hrs* knockdown efficiency in AML12 cells used in (**A**). **C.** Confocal micrographs showing ORO-stained LDs (red) and DAPI-stained nuclei (blue) in AML12 cells transfected with control (siNT) or *Hrs*-targeting (si*Hrs*) siRNAs, followed by a pulse treatment of OA and a chase period in growth medium without OA (WD). **D.** Quantification of total LD Area per cell in cells from (**C**). **E.** Immunoblot analysis confirming *Hrs* knockdown efficiency in AML12 cells used in (**C and D**). **F.** Quantification of total LD area per cell in AML12 cells transfected with control (siNT) or *Hrs*-targeting (si*Hrs*) siRNAs subjected to a pulse treatment of OA followed by a chase period in growth media (WD) with or without ATGLi. **G.** Immunoblot analysis confirming *Hrs* knockdown efficiency in AML12 cells used in (**F**). Data represent the mean ± SD from three independent trials, where each dot represents an independent trial. The statistical significance is indicated by asterisks using a two-way ANOVA with a (**A**) Sikak’s, (**D**) Uncorrected Fisher’s LSD, or a (**F**) Tukey’s post-hoc test, \**P* < 0.05, \*\**P* < 0.01, \*\*\**P* < 0.001, \*\*\*\**P* < 0.0001, n.s., not significant.

### HRS regulates LD catabolism independent of lipolysis

To assess if HRS affects LD catabolism, siNT and si*Hrs* AML12 cells were first pulse-loaded with 150 µM oleic acid for 4 h to stimulate LD biogenesis, then washed and chased in regular medium without OA for an additional for 24 h. Following this 24 h withdrawal period (24 h WD in **Figure 2C-E; Figure S2C, D**), siNT cells had a significant reduction in LD area/cell (∼ 50 %). However, cells lacking HRS had ∼ 25 % reduction in LD area/cell following the 24 h withdrawal period (**Figure 2C-E**), suggesting that LD catabolism occurs at a reduced rate in HRS-deficient cells.

LD catabolism occurs by lipolysis and lipophagy pathways. In hepatocytes, accumulation of large LDs indicates a defect in lipolysis, whereas small LDs indicate a defect in lipophagy.^2^ Patatin-like phospholipase domain-containing protein 2 (PNPLA2, commonly known as ATGL) is a well-characterized enzyme that catalyzes the rate-limiting step of lipolysis. If lipolysis is perturbed by *Hrs* knockdown, then *Hrs* knockdown cells would exhibit reduced sensitivity to the ATGL inhibitor, Atglistatin (ATGLi). We performed the LD catabolism experiment described above, where we loaded the siNT and si*Hrs* cells with 150 µM OA, followed by a 24 h OA withdrawal period (indicated as “WD” in **Figure 2F; Figure S2E, F**) with or without ATGLi. As depicted in **Figure 2F**, ATGLi treatment of siNT cells during the 24 h WD period resulted in a significant increase in LD area/cell (∼ 2.5-fold) compared to the 24 h WD period without ATGLi. Further, ATGLi treatment of si*Hrs* resulted in a modest increase (∼ 1.5-fold) in LD area/cell (**Figure 2F, G**), without affecting LD number/cell or LD size (**Figure S2E, F**, respectively), compared to the 24 h WD period without ATGLi. The observed increase in LD area/cell levels suggests *Hrs* knockdown does not significantly hinder lipolysis activity. This is consistent with the observation that *Hrs* knockdown does not affect ATGL protein levels (**Figure S2G, H**). Altogether, the accumulation of LDs observed in HRS-deficient cells is not solely due to defects in lipolysis.

### HRS knockdown affects both macro– and micro-lipophagy pathways

Since ESCRT proteins were found to physically interact with LDs and regulate LD trafficking to the yeast lysosomes (vacuoles),^10^ we first analyzed whether HRS co-localized with LDs by using bioinformatic and proteomic analyses provided by the lipid droplet knowledge portal.^21, 22^ This portal contains data from an unbiased proteomics screen assessing mouse liver protein-organellar localization by isolating and enriching for cellular organelles by sucrose gradient.^22^ Using this database, HRS protein from mouse liver was not present in cellular isolations enriched for LDs (**Figure S3A**). Instead, cellular isolations containing HRS overlapped with those containing endosomal marker proteins (**Figure S3A**). We confirmed these results through fluorescence microscopy in AML12 cells, which showed negligible co-localization between endogenous HRS and LDs (**Figure S3B**) and even upon HRS overexpression (**Figure S3C**). Thus, we concluded that HRS regulates LD degradation independent of LD localization. Next, we sought to test the role of HRS in lipophagy (i.e. LD trafficking to the lysosome) using AML12 cells subjected to a “lipophagy flux” pulse-chase assay as previously described in Schott et al.^4^ In brief, siNT and si*Hrs* cells were pulse-loaded with OA (150 µM) and fluorescent BODIPY 558/568 C_12_ (7.5 µM) for 4 h to label LDs, then washed and chased in DMSO vs LAL inhibitor (LAListat or LALi, 10 µM) for an additional 24 h (**Figure 3A**).^4^ We predicted that if HRS facilitates LD-lysosome trafficking, then LD colocalization with lysosomes would be decreased in *Hrs* knockdown cells, particularly when treated with LALi. Surprisingly, si*Hrs* cells displayed enlarged LAMP1-GFP vesicles with the accumulation of numerous LDs within the lysosomal lumen in both DMSO and LAListat-treated cells (**Figure 3B-D**). This suggests *Hrs* knockdown does not prevent LD-lysosome trafficking but may cause lysosome defects leading to a redued capacity to degrade LDs. Furthermore, higher instances of LD-lysosome colocalization events within si*Hrs* cells suggests that LD-lysosome trafficking may be accelerated.

**Figure 3.**
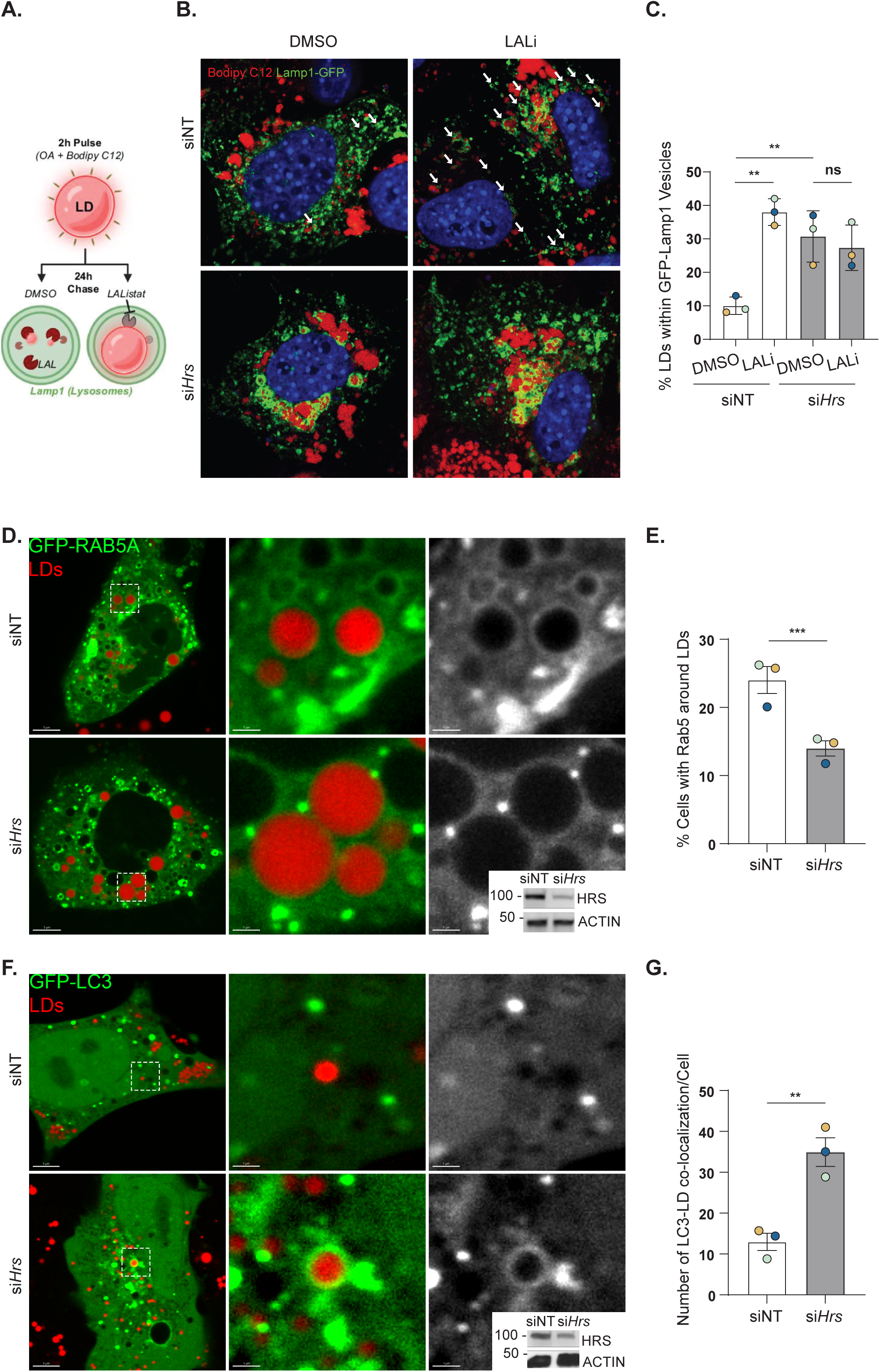
HRS knockdown affects both micro– and macro-lipophagy pathways. **A.** Schematic illustrating pulse-chase assay to observe LDs using fluorescent BODIPY 558/568 C12 within LAMP1-GFP vesicles (lysosomes). **B.** Confocal micrographs showing AML12 cells transfected with control (siNT) or *Hrs*-targeting (si*Hrs*) siRNAs and Lamp1-GFP (green), BODIPY-stained LDs (red), DAPI-stained nuclei (blue), with or without LAListat treatment. White arrows indicate LDs within Lamp1-GFP-positive vesicles. **C.** Quantification of the percentage of LDs within Lamp1-GFP-positive vesicles in cells from (**B**). **D.** Immunoblot analysis confirming *Hrs* knockdown efficiency in AML12 cells used in (**B,C**). **E.** Confocal micrographs showing AML12 cells transfected with control (siNT) or *Hrs*-targeting (si*Hrs*) siRNAs and GFP-tagged RAB5 (green). LDs were stained with MDH (red). **F.** Quantification of the percentage of cells with RAB5 around LDs in (**E**). **G.** Immunoblot analysis confirming *Hrs* knockdown efficiency in AML12 cells used in (**E, F**). **H.** Confocal micrographs showing AML12 cells transfected with control (siNT) or *Hrs*-targeting (si*Hrs*) siRNAs and GFP-tagged Map1lc3 (green). LDs were stained with MDH (red). **I.** Quantification of the percentage of cells with LC3 around LDs IN (**H**). **J.** Immunoblot analysis confirming *Hrs* knockdown efficiency in AML12 cells used in (**H,I**). Data represent the mean +/− SD from three independent trials, where each dot represents an independent trial. The statistical significance is indicated by asterisks using a (**C**) one-way ANOVA with a Tukey’s post-hoc test, or (**F, I**) a two-tailed Student’s t-test for ratio-pairwise comparisons, \*\**P* < 0.01, \*\*\**P* < 0.001, n.s. not significant.

To determine if HRS affects microlipophagy in AML12 cells, we probed the localization of the early endosome protein RAB5, which has been previously shown to facilitate LD-lysosome trafficking by microlipoophagy.^4^ As seen in Figure 3E-G, si*Hrs* cells exhibited a ∼ 50 % reduction in GFP-RAB5 targeting to LDs, indicating impaired targeting by microlipophagy. This observation was curious given the extreme accumulation of LDs within lysosomes (**Figure 3B, C**), suggesting perhaps a compensatory mechanism that overcomes the defective microlipophagy. To test this, we probed the impact of *Hrs* knockdown on macrolipophagy by analyzing LC3-positive autophagosomes near LDs using confocal microscopy. As seen in **Figure 3H-J**, LC3 targeting to LDs was increased ∼ 3-fold in HRS-depleted AML12 cells compared with siNT, indicating increased LD targeting by macrolipophagy machinery. Altogether, these results suggest that HRS knockdown diminishes Rab5-mediated macrolipophagy, while causing a simultaneous increase in autophagosome-dependent trafficking of LDs to lysosomes.

### Autophagy flux is perturbed in HRS-depleted cells

To test whether HRS depletion had a broader effect on autophagic flux, siNT and si*Hrs* AML12 cells were treated with ± mTOR inhibitor Torin1 (250 nM, 30 min) to stimulate autophagy, and ± lysosomal inhibitor bafilomycin A1 (BafA1; 100 nM, 2 h) to measure the accumulation of autophagic cargo. The autophagosome marker LC3-II and the autophagic cargo protein SQSTM1 were assessed by western blot analysis (**Figure 4A**). As shown in **Figure 4B, C**, under basal conditions (without Torin1) and stimulated autophagy (with Torin1), BafA1 caused a substantial increase in LC3-II accumulation in non-targeting control cells (Basal, 15-fold increase; Stimulated, 3.5-fold increase), but this accumulation was consistently blunted in HRS-depleted cells (Basal, 5-fold; Stimulated, 1.9-fold increase by BafA1). Analysis of BafA1-mediated accumulation of LC3-II showed a significant reduction in autophagic flux in HRS-depleted cells compared with non-targeting control cells (**Figure 4D**). Interestingly, LC3-II levels were higher in HRS-depleted cells (without BafA1), suggesting that autophagosome formation may initially be enhanced without HRS, while lysosomal catabolism may be simultaneously decreased. Analysis of p62/SQSTM1 initially suggested that autophagic cargo degradation may be enhanced in HRS-depleted cells (**Figure 4E**), but this effect was not observed under autophagy-stimulating conditions using Torin1 (**Figure 4F-G**). Taken together, these results suggest that autophagic flux is reduced in HRS-depleted AML12 cells.

**Figure 4.**
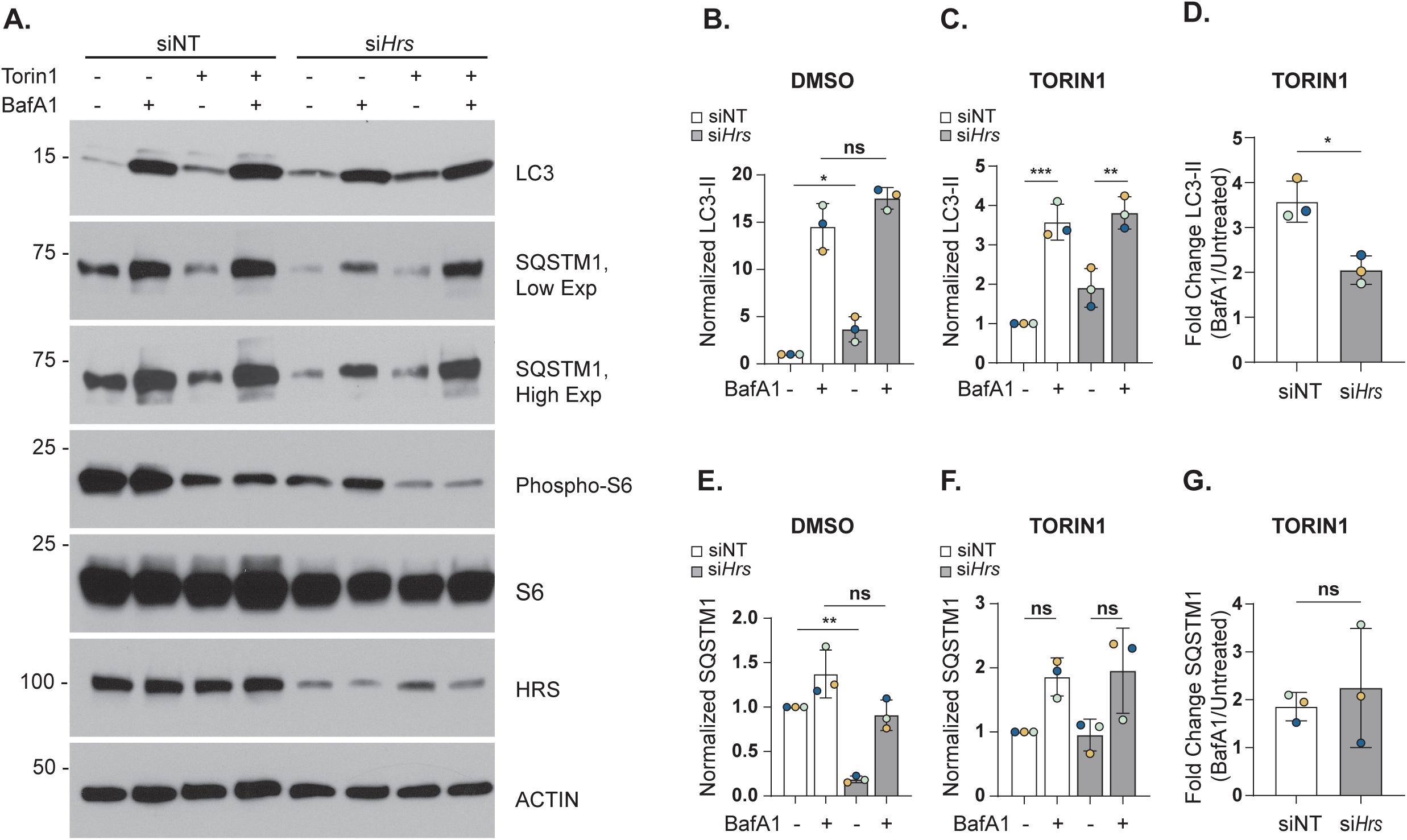
Autophagy flux is decreased in HRS-deficient cells. **A.** Representative immunoblot autophagic flux in AML12 cells transfected with control (siNT) or *Hrs*-targeting (si*Hrs*) siRNAs and treated with or without BafA1 followed by DMSO or Torin1. Quantification of LC3-II in siNT or si*Hrs* cells with (**B**) DMSO or (**C**) Torin1 and ± BafA1. Changes in LC3-II/ACTIN for each group was normalized against siNT. **D.** Quantification of fold change in LC3-II ± BafA1 from Torin1-treated cells as a measure of autophagic flux. **E.** Quantification of SQSTM1 in siNT or si*Hrs* cells with DMSO or (**F**) Torin1 ± BafA1. **G.** Quantification of fold change in SQSTM1 levels in Torin1-treated cells ± BafA1. Data represent the mean ± SD from three independent trials, where each dot represents an independent trial. The statistical significance is indicated by asterisks using a (**B, C, E, F**) one-way ANOVA with Tukey’s post-hoc test, or (**D, G**) two-tailed Student’s t-test for ratio-pairwise comparisons, \**P* < 0.05, \*\**P* < 0.01, \*\*\**P* < 0.001, n.s., not significant.

### HRS deficiency inhibits mTOR signaling and activation

In the autophagic flux experiments above, we also analyzed p70 ribosomal S6 protein phosphorylation, a downstream substrate of the mTOR pathway, to confirm that Torin1 was indeed decreasing mTOR activity as expected. Interestingly, we also observed that HRS depletion itself seemed to dramatically reduce mTOR activity similar to Torin1 treatment. To test this, we expanded our analysis of mTOR phosphorylation targets to include ULK1, p70 ribosomal S6 kinase (S6K), and 4E-BP1.^23^

As shown in Figure 5, *Hrs* knockdown in AML12 cells caused a ∼ 50% reduction in phosphorylation of all mTOR substrates examined (**Figure 5A-E**). As expected, Torin1 treatment significantly reduced phosphorylation of these proteins in control cells (**Figure 5A-E**). Additionally, total protein levels of ULK1, 4E-BP1, and S6K were reduced by more than half in si*Hrs* cells, while S6 levels remained unaffected (**Figure S4A-D**). These findings support the conclusion that HRS depletion leads to inhibition of mTOR activity.

**Figure 5.**
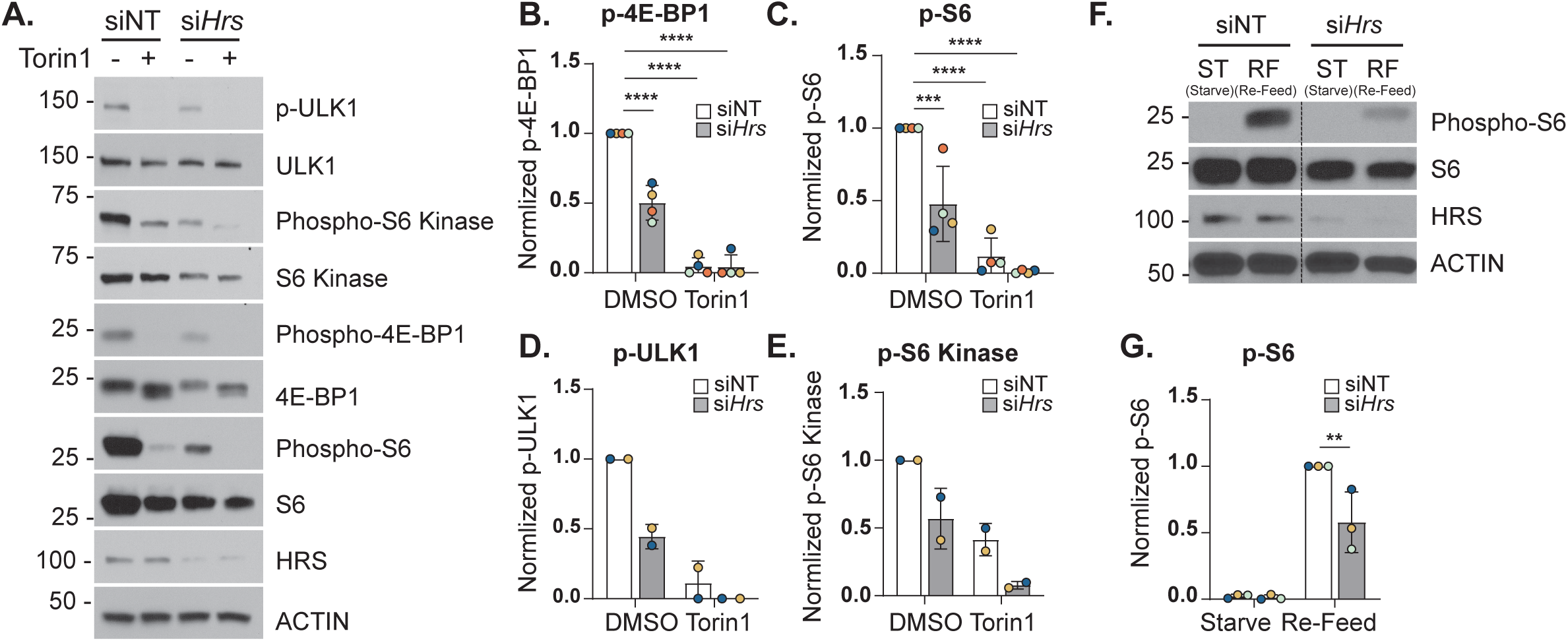
mTOR signaling is decreased in HRS-deficient cells. **A.** Representative immunoblot showing mTOR activity in siNT and si*Hrs* cells treated ± Torin1. Quantification of (**B**) phospho-4E-BP1, (**C**) phospho-S6, (**D**) phospho-ULK1, (**E**) phospho-S6 kinase in siNT or si*Hrs* cells treated ± Torin1. Each phospho-protein was normalized to their respective total protein. **F**. Representative immunoblot showing nutrient-stimulated mTOR activity in siNT and si*Hrs* cells upon starvation in HBSS media (ST) before re-feeding with complete media (RF). **G.** Quantification of phospho-S6 in siNT or si*Hrs* cells normalized to total S6. Data represent the mean ± SD from three independent trials, where each dot represents an independent trial. The statistical significance is indicated by asterisks using a two-way ANOVA with (**B-E**) Dunnett’s, or (**G**) Uncorrected Fisher’s LSD post-hoc test, \*\**P* < 0.01, \*\*\**P* < 0.001, \*\*\*\**P* < 0.0001, n.s., not significant.

To corroborate these findings, we evaluated the impact of HRS depletion on stimulated mTOR activity following starvation and subsequent nutrient refeeding. siNT and si*Hrs* AML12 cells were subjected to nutrient deprivation using Hank’s Balanced Salt Solution (HBSS) for 1 h, then refeeding with nutrient-rich medium for 10 min as previously described.^24^ As expected, starvation abolished TORC1 signaling in both groups, evidenced by loss of phosphorylated S6 (indicated as “ST” in **Figure 5F**). Upon refeeding, control cells exhibited robust TORC1 reactivation (indicated as “RF” in **Figure 5F**), whereas HRS-deficient cells displayed more than a 50 % reduction in recovery compared with controls (**Figure 5F, G**). These results establish HRS as a positive regulator of mTOR signaling and are consistent with the above findings that HRS depletion could increase autophagosome formation (**Figure 4A, B**) despite an overall decrease in autophagic flux which could be attributed to defective lysosomes (**Figure 3**).

### HRS does not affect lysosomal levels but regulates lysosomal activity

Because HRS-deficient cells can still traffic LDs to lysosomes – likely through increased autophagosome targeting – but retain LDs within lysosomes similar to that of LAListat-treated cells (**Figure 3**), we next asked whether this accumulation reflects impaired lysosomal degradation. First, Western blot analysis of LAMP1 levels revealed no significant differences between siNT and si*Hrs* AML12 cells (**Figure 6A, B**), suggesting lysosome abundance is not affected by HRS depletion.

**Figure 6.**
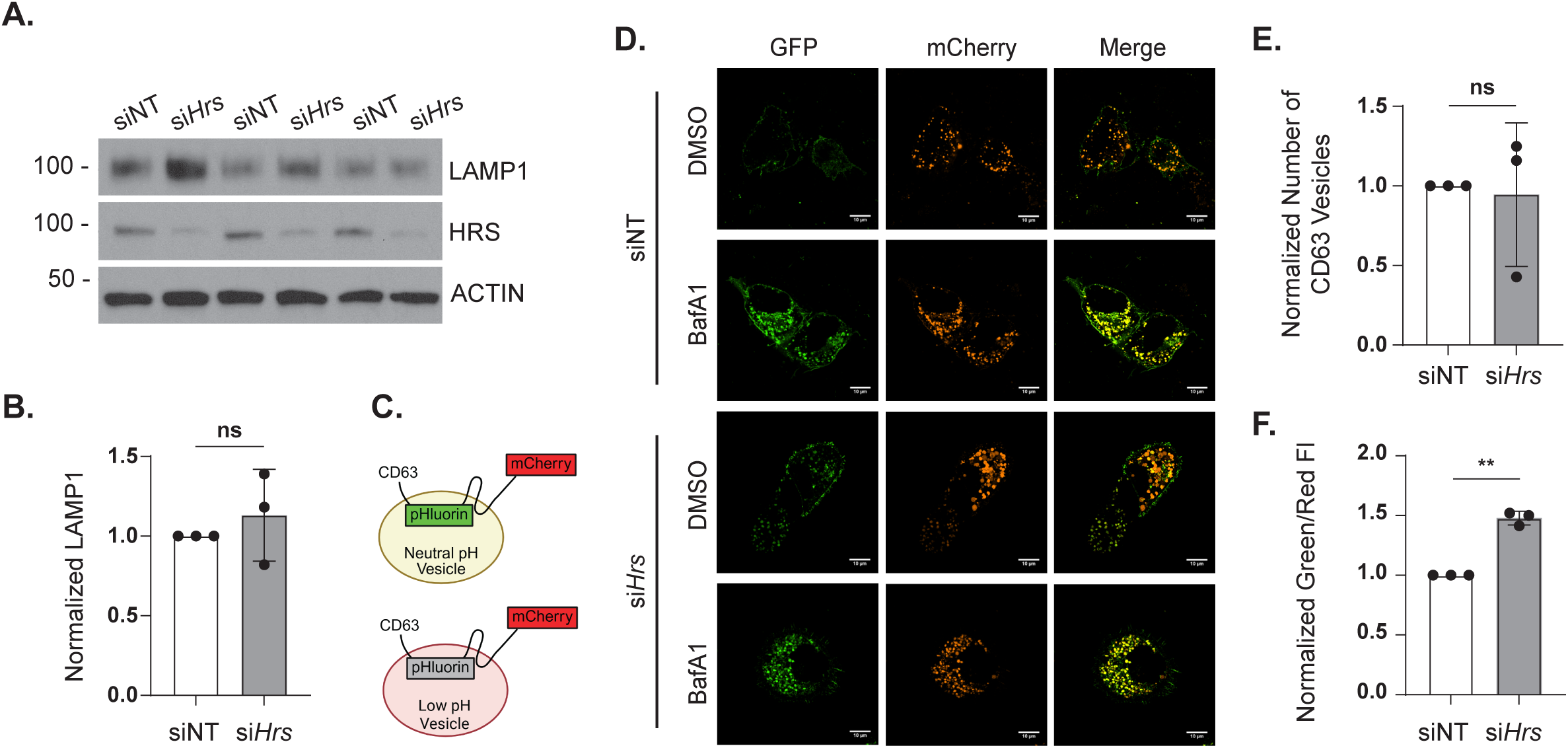
HRS regulates lysosomal function. **A**. Representative immunoblot and (**B**) quantification of LAMP1 levels in siNT and si*Hrs* AML12 cells. **C.** Schematic illustrating CD63-pHluorin-mCherry pH reporter construct. **D.** Confocal micrographs of siNT and si*Hrs* AML12 cells transfected with CD63-pHluorin-mCherry reporter and treated with BafA1 or DMSO. **E.** Quantification of CD63 vesicles per cell and (**F**) green/red fluorescence intensity ratio in siNT and si*Hrs* AML12 cells. Data represent the mean ± SD from three independent trials, where each dot represents an independent trial. The statistical significance is indicated by asterisks using a two-tailed Student’s t-test for ratio-pairwise comparisons. ns = not significant, \*\**P* < 0.01.

We next assessed lysosomal acidity using a dual-fluorescence CD63-pHluorin-mCherry reporter, which localizes to lysosomes and late endosomes. The pH-sensitive pHluorin localizes to the vesicle lumen, where its fluorescence is quenched under acidic conditions (**Figure 6C**).^25^ mCherry fluorescence is pH-insensitive and serves as an internal control for intracellular expression levels. In this system, elevations in lysosome pH correlate with higher green/red fluorescence ratios and appear as yellow puncta. siNT and si*Hrs* cells expressing the reporter were imaged live with or without treatment with BafA1. While HRS depletion did not alter the number of CD63-positive vesicles (**Figure 6E**), the ratio of green/red fluorescence was increased by 1.5-fold compared with siNT cells, indicating perturbed lysosomal acidity (**Figure 6D, F**). Together, these findings demonstrate that HRS deficiency perturbs lysosomal pH and thereby compromises lysosomal degradative function. This impairment likely contributes to the accumulation of LDs within lysosomes (**Figure 3**), and decreased autophagic flux (**Figure 4**) observed in HRS-deficient cells.

## Discussion

Yeast studies, where microlipophagy was first described, have implicated the ESCRT proteins in facilitating LD trafficking to the yeast vacuole. Later studies have found the ESCRT proteins Chmp1 and Chmp4a to regulate LDs in flies,^15^ while VPS4A and VPS4B were found to regulate LDs in hepatocytes.^14^ Our unbiased screen of ESCRTs and ESCRT adaptors identified several proteins that regulate LDs in Hep3B and HeLa cells (**Figure 1**). Overall, this growing body of literature demonstrates the conserved role of ESCRT proteins in regulating lipophagy from simple eukaryotes to complex ones. Interestingly, while ESCRT proteins were identified to regulate lysosomal pore closure after LD internalization in yeast^26^, they appear to have gained different functions over the course of evolution, acting as lipophagy receptors with LC3-interacting regions (LIRs) in the case of VPS4A^14^ and regulating lysosome pH, as shown by our data.

Our results show that knockdown of the ESCRT-0 protein HRS caused a significant increase in LD levels in Hep3B, HeLa, HepG2, and AML12 cells (**Figure 1 and S1**). Studies so far have only implicated HRS in recruiting subsequent ESCRT sub-groups and coordinating the transport of ubiquitinated cargo for lysosomal degradation.^16–18^ This role in lysosomal degradation of ubiquitinated cargo makes HRS an important regulator of autophagy pathways involved in protein degradation across cell types. As examples, HRS plays a role in regulating the degradation of ubiquitinated and phosphorylated receptors on endosomes through a process called simaphagy,^12^ prevents protein aggregate-mediated ER stress,^27^ and regulates myogenesis through the positive regulation of the AKT signaling pathway and the negative regulation of MEK/ERK signaling pathway.^28^ Additionally, HRS plays a role in lysosomal degradation of other biomolecules, including epidermal growth factor receptor (EGFR), leading to receptor accumulation within early endosomes and prolonged activation.^29, 30^ Overall, HRS has functional relevance in multiple lysosomal degradation pathways in the cell, including lipophagy.

Seminal research from Oku, et al. has revealed the yeast ESCRT protein Vps27 (i.e. the yeast ortholog of HRS) as a regulator of microautophagy and microlipophagy following a metabolic shift where the carbon source is changed from glucose to ethanol (i.e. diauxic shift).^10^ Vps27-dependent microautophagy was found to be dependent on all of its canonical functions (i.e., binding to ubiquitin, PI3P, and ESCRT-I factors). To evaluate microlipophagy, Oku, et al. assessed cleavage of fluorescent proteins fused with the LD resident protein, Osw5 (Osw5-EGFP). After assessing cleavage of Osw5-EGFP, they found that 1) its cleavage is dependent on intravacuolar protease activity and 2) loss of Vps27 caused significantly reduced cleavage rates. Moreover, they also concluded that the conserved clathrin-binding motif (also present in mammalian HRS) of Vps27 is indispensable for cleavage, where yeast expressing Vps27 mutants lacking this motif had fewer LDs within the vacuole (i.e., LD-vacuole trafficking).^10^ Similarly, our RNAi screen identified HRS as a regulator of lipophagy in mammalian cells, however, our *Hrs* knockdown cells did not result in fewer LDs within lysosomes (**Figure 3**). While we did not test different HRS mutants, it would be interesting to evaluate if and how the HRS’s ubiquitin– and clathrin-binding motifs of HRS would affect lipophagy in mammalian cells.

LDs from HRS-deficient cells drastically accumulate in the lysosome, where lysosomal lipase inhibition does not result in an additional increase in LDs within this compartment (**Figure 3**). To further elucidate the LD-lysosome trafficking mechanism in HRS-deficient cells, we probed micro– and macro-lipophagy by assessing the colocalization of LDs with RAB5 (a known regulator of microlipophagy in hepatocytes)^4^ and LC3, respectively. We noticed a significant decrease in LD targeting by RAB5 and a profound increase in LD-autophagosome targeting in HRS-deficient cells (**Figure 3**). These results support the hypothesis that HRS is necessary for LD targeting by RAB5.

The observed upregulated macrolipophagy and extreme accumulation of LDs within lysosomes prompted us to assess general aspects of autophagy, e.g. autophagosome formation and degradation in the lysosome. Our assays reveal that the autophagic flux in HRS-deficient cells is significantly diminished (**Figure 4**). This is consistent with a previous report that found autophagy flux in neurons to be blocked considerably upon *Hrs* knockdown.^27^ Notably, HRS regulates the recruitment and sorting of membrane receptors (e.g., EGFR) into multi-vesicular bodies. facilitating their trafficking to the lysosome for degradation.^27^ Interestingly, HRS was found to modulate the recruitment of autophagic regulators like SQSTM1 and NBR1 to endosomes potentially affecting their availability to participate in autophagic processes.^12^ On the other hand, the increased targeting of LDs by autophagosomes is an interesting phenomenon, as such compensatory dynamics of macro– and micro-lipophagy pathways have not been characterized previously. Future studies will be necessary to discern whether these results occur specifically from HRS activity or are due to more broad-ranging defects in endosomal trafficking that occur from HRS depletion. Overall, our results support the role of ESCRT proteins in autophagy. Many ESCRT proteins play critical roles during different stages of autophagy, from autophagosome formation to their fusion with lysosomes.^31, 32^ Thus, our results suggest a role for HRS in autophagy flux regulation that aligns with the current literature. Our autophagy flux experiments revealed that *Hrs* knockdown cells had decreased phosphorylated S6 protein (**Figure 5**). S6 protein is a downstream target of mTOR, a master inhibitor of general autophagy.^23^ We examined phosphorylation levels of other mTOR targets to see if inhibition was limited to S6 or affected other mTOR targets. Surprisingly, HRS acts as a positive regulator of mTOR signaling. *Hrs* knockdown cells show decreased phosphorylation of mTOR targets, like treatment with the mTOR inhibitor Torin1 (**Figure 5**). Considering autophagy flux is decreased in *Hrs* knockdown cells, this result was perplexing as decreased mTOR signaling should result in increased autophagy. However, a similar phenomenon has been observed in podocytes where decreased mTOR signaling caused accumulation of the autophagosome marker LC3 (possibly indicating decreased autophagy flux).^33^ Also, the effect of HRS on mTOR could be similar to the mechanism of ESCRT-1-dependent mTOR regulation, where the depletion of ESCRT-I proteins TSG101 and VPS28 reduced RAG-GTPase-dependent mTOR signaling.^34^ Furthermore, mTOR signaling inhibition also leads to defects in lysosomal activity.^35^ Thus, we hypothesized that LD accumulation and decreased autophagy flux could be due to defective lysosomes.

Our results show that *Hrs* knockdown does not affect LAMP1 levels (**Figure 6A**), indicating lysosomal abundance is not affected. Alternatively, defects in lysosomal degradation could also be due to an increase in lysosomal pH that renders the lysosomes non-functional.^36^ We found that lysosome pH is significantly increased in *Hrs* knockdown cells (**Figure 6D, F**). This lysosomal defect is consistent with the observed LD accumulation within lysosomes in HRS-depleted cells.Thus, the effect of HRS on lysosomal pH could indirectly affect mTOR and/or mTORC1.

In conclusion, our results further support the idea that ESCRT proteins regulate LD levels in multiple mammalian cells, including hepatocytes. In addition to its canonical functions, we identified that HRS has unprecedented roles in regulating lipophagy. It affects micro– and macro-lipophagy pathways, general autophagic flux, and maintains lysosome function (**Figure 7**). These findings are significant, especially in the context of lipid trafficking and utilization disorders, including steatohepatitis, where hepatocytes accumulate LDs and could potentially cause irreversible liver damage. Thus, elucidating ESCRT-mediated lipophagy regulation in mammalian cells can potentially aid in developing novel targets to ameliorate fat accumulation diseases of the liver.

**Figure 7.**
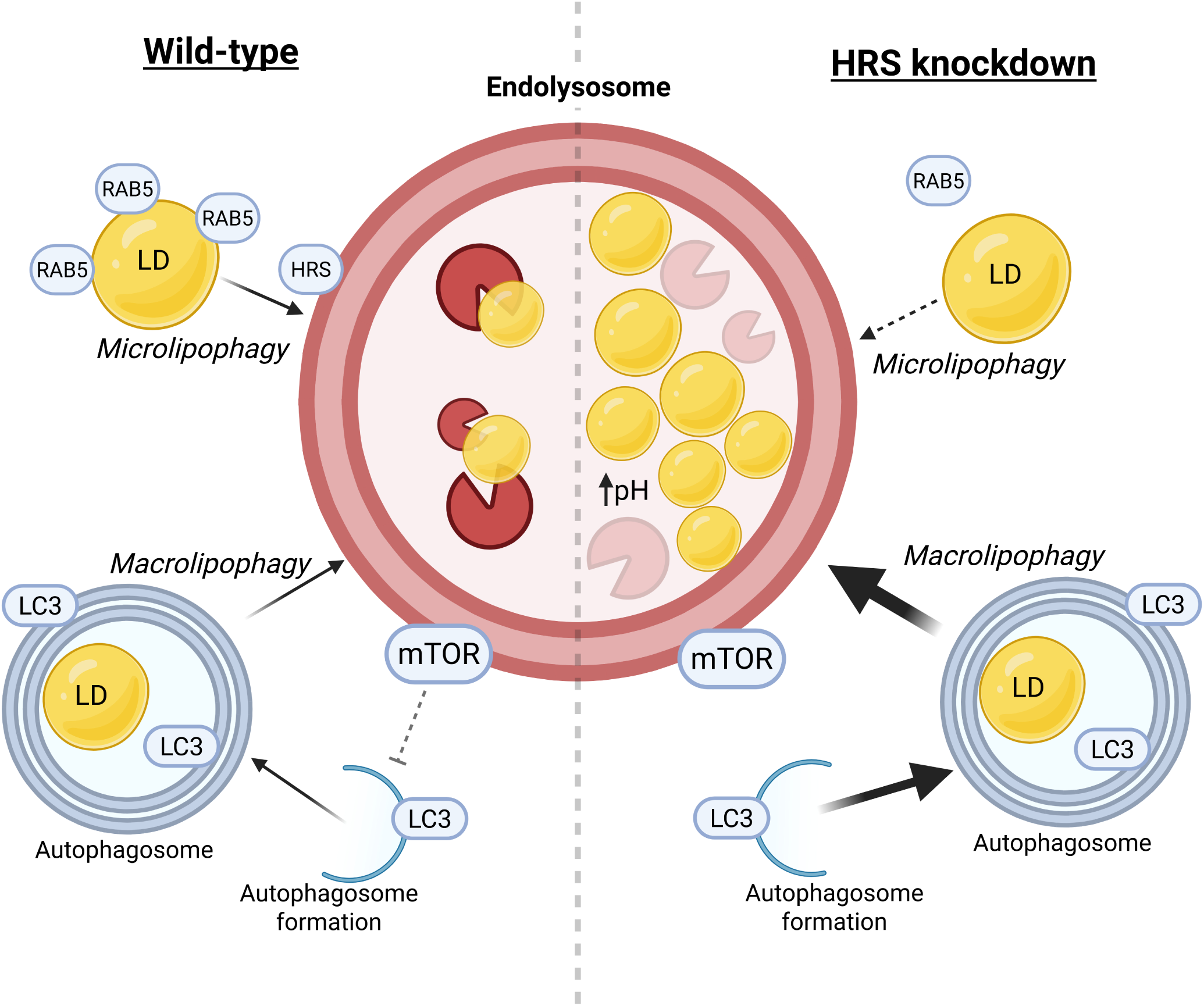
Model depicting HRS-mediated regulation of lipophagy and lysosomal function. HRS positively regulates RAB5-targeting of LDs during lipophagy. Alternatively, HRS negatively regulates autophagosome-targeting of LDs for lipophagy. Further, HRS facilitates lysosomal function and general autophagic degradation by maintaining acidic conditions. The observed increase in autophagosome-dependent targeting of LDs in combination with dysfunctional lysosomes in HRS-deficient cells causes a significant accumulation of LDs within lysosomes.

## Materials and Methods

### Cell Lines, Media Component, and Immunological Reagents

Hep3B (ATCC HB-8064) cells were maintained in Eagle’s Minimum Essential Medium (MEM) (FisherScientific MT10009CV) supplemented with 10 % FBS (Gibco 26140079), 1 % Pen/Stre (FisherScientific 15140-122) at 5 % CO_2_, 37 °C. HeLa (ATCC CCL-2) cells were maintained in DMEM (FisherScientific MT10013CV) supplemented with 10 % FBS, 1 % Pen/Strep. AML12 (ATCC CRL-2254) cells were maintained in DMEM/F-12 1:1 media (ThermoFisher 11330057) supplemented with 10% FBS (Gibco 26140079), 1% Pen/Strep (Fisher Scientific 15140-122), and 1X ITS (Gibco 41400045) at 5 % CO_2_, 37 °C. Trypsin (Fisher Scientific 15-400-054) diluted in HBSS w/o Ca, Mg (Fisher Scientific MT21021CV) was used to passage cells.

### Screening Analysis

Hep3B and HeLa cells were grown in duplicate wells of a glass-bottom 96-well plate (Ibidi, n=3 independent RNAi experiments). Following 72 h of knockdown, cells were stained with Hoechst and LipidTOX green and imaged using high-throughput epifluorescence microscopy. Total LD area/cell was quantified using CellProfiler image analysis.

### siRNA Knockdown

AML12 cells were grown to 30 % – 50 % confluency before transfection using Lipofectamine RNAiMAX (Thermo Fisher 13778-150). All transfections were done using manufacturer-recommended conditions. siOnTARGET pools targeting mouse *Hrs* (Horizon Discovery L-055516-01-0005) or a nontargeting control siRNA (Horizon Discovery D-001810-10-20) were used for 48 h knockdowns. Knockdowns were confirmed for each replicate of all assays by immunoblotting.

### Oil red O staining

The Oil Red O (ORO) staining was performed by fixing AML12 cells with 4 % paraformaldehyde (Tousimis Research Corporation #1008D). Fixed cells were washed with 60 % 2-propanol for 30s, then stained with ORO (Sigma O0625) for 90 s. This was followed by a second wash with 60 % 2-propanol for three quick (∼ 30 s) d-PBS washes. The coverslips were mounted on slides with Prolong with DAPI (for counterstaining the nucleus, ThermoFisher P36931) and imaged using a Nikon Eclipse Ti2 microscope. The stain-positive region of each image were analyzed using ImageJ (Auto Local Threshold tool, Bernsen method), previously used in Schott, et al.^2^

### Western Blotting

AML12 cells grown in either 6-well plates or 10 cm dishes and were silenced for 48 hours. For assays with further treatments, an additional well or dish with the same number of AML12 cells was transfected with control and Hrs-targeting siRNA. Cells were lysed in 1X RIPA buffer containing 1 Complete Mini, EDTA-free, Protease Inhibitor Cocktail Tablet (Sigma 11836170001), and Halt phosphatase inhibitor (VWR PI78427), and sonicated (Fisher Scientific FB120) using Amp 70 %, pulsing 3x for 3 s each with 2 s in between. Post-nuclear supernatant (PNS) was collected following centrifugation at 16,100 x g (Eppendorf 5415R) for 5 min at 4 °C. Total protein concentrations were determined using the Pierce BCA protein assay kit (ThermoFisher #23227) per the manufacturer’s recommended protocols. Loading samples were prepared in 4X Laemli sample buffer (Biorad 1610747) boiled at 100 °C for 5 minutes. Samples were loaded into 10 % or gradient 4-20% tris-glycine gels (BioRad 4561034 and 4561094, respectively) for SDS-PAGE. Following SDS-PAGE proteins were transferred to polyvinylidene difluoride (PVDF) membranes (Sigma IPVH00010) using a semi-dry transfer apparatus (IBI IB45000) for 30 minutes. Membranes were processed with the desired primary and secondary antibodies per the manufacturer’s recommended protocol and concentrations. List of antibodies used in the study is included in Table 1. Immunolabeled bands were detected by chemiluminescence using the ECL substrate (ThermoFisher 34580) and autoradiography film (MidSci BX810) or imaged on a C-digit imaging system (LiCor Biotech). Densitometric analysis of the bands was performed using ImageJ (National Institutes of Health, Bethesda, MD).

**Table 1:** List of antibodies.

| **Target protein** | **Antibody type** | **Manufacturer, Catalog #** |
| --- | --- | --- |
| 4E-BP1 | Primary antibody | Cell Signaling Technologies, 9644S |
| ACTIN | Primary antibody | Sigma Aldrich, A2066-2ML |
| HRS | Primary antibody | Cell Signaling Technologies, 15087S |
| LC3B | Primary antibody | Cell Signaling Technologies, 2775S |
| Phospho-4E-BP1<br>(T37/46) | Primary antibody | Cell Signaling Technologies, 2855S |
| Phospho-S6 (S240/244) | Primary antibody | Cell Signaling Technologies, 5364 |
| Phospho-S6 Kinase<br>(S371) | Primary antibody | Cell Signaling Technologies, 9208T |
| Phospho-ULK1 | Primary antibody | Cell Signaling Technologies, 6888S |
| S6 | Primary antibody | Cell Signaling Technologies, 2217 |
| S6 Kinase | Primary antibody | Cell Signaling Technologies, 34475S |
| SQSTM1/p62 | Primary antibody | Cell Signaling Technologies, 5114S |
| ULK1 | Primary antibody | Cell Signaling Technologies, 8054S |
| LAMP1 | Primary antibody | Cell Signaling Technologies, 9091S |
| ATGL (PNPLA2) | Primary antibody | Cell Signaling Technologies, 2439S |
| Anti-Rabbit IgG HRP | Secondary<br>antibody | Cell Signaling Technologies, 7074P2 |

### Lysosome acidity

The CD63-pHlourin-mCherry plasmid was a gift from Dr. David Katzmann (Mayo Clinic). After transfecting AML12 cells with siRNAs for 48 hours, cells were transfected with the CD63-pHlourin-mCherry reporter plasmid using Lipofectamine 2000 (Fisher Scientific 11668-027) for 24 hours. Live cell imaging was performed on an Andor BC43 microscope from Oxford Instruments and on a Nikon Eclipse Ti2 microscope equipped with an NSPARC detector. The images from Andor BC43 microscope were analyzed for red and green fluorescence intensity of each cell and were measured using ImageJ. The images taken on a Nikon Eclipse Ti2 microscope were used for the representative images. Protein lysates from control and Hrs-knockdown cells were analyzed for LAMP1 expression apart from HRS and housekeeping protein ACTIN.

## Acknowledgements

This work was supported by National Institutes of Health grants R35GM150801 (M.B.S) and R00AA026877 (M.B.S).

The content is solely the responsibility of the authors and does not necessarily represent the official views of the National Institutes of Health. The authors declare no competing financial interests.

Author contributions: M.M. Willoughby, A. Shroff, and M.B. Schott conceived the study, designed experiments, interpreted data, and wrote the manuscript. M.M. Willoughby, A. Shroff, B.E. Crossman, and M.B. Schott performed and analyzed experiments.

## Abbreviations

LD: Lipid Droplet
HRS: hepatocyte growth factor receptor tyrosine kinase substrate
LAL: Lysosomal Acid Lipase
ESCRT: Endosomal Sorting Complexes Required for Transport
BafA1: Bafilomycin A1

## Supplemental Figures

**Figure S1.** **A**. Graph representing fold changes in total LD area per cell in Hep3B cells post knockdown of indicated ESCRT genes. **B.** Graph showing changes in total LD area per cell from RNAi screen of ESCRT and ESCRT-related genes in HeLa cells. **C.** Confocal micrographs of AML12 cells transfected with *Hrs*-targeting siRNA for 48 hours. The cells were then analyzed by immunofluorescence for early endosomes (EEA1, red), HRS (green), and DAPI-stained nuclei (blue). Quantification of total LD area per cell in (**D**) HeLa and (**E**) HepG2 cells treated with control (siNT) versus *Hrs* targeting (si*Hrs*) siRNAs for 48 hours. Data represents the mean ± SD from at least three independent trials, where each dot represents an independent trial. The statistical significance is indicated by asterisks using a two-tailed Student’s T-test for ratio-pairwise comparisons. *\*P < 0.05, **P < 0.01*, n.s., not significant. Abbreviations: ESCRT – Endosomal Sorting Complex Required for Transport, LD: Lipid Droplet.

**Figure S2.** Quantification of (**A**) LD number per cell and (**B**) LD size in AML12 cells transfected with control (siNT) or *Hrs*-targeting (si*Hrs*) siRNAs and treated with OA, with or without DGAT1/2i. Quantification of (**C**) LD number per cell and (**D**) LD size in AML12 cells transfected with control (siNT) or *Hrs*-targeting (si*Hrs*) siRNAs subjected to a pulse treatment of OA followed by a chase period in growth media without OA for 24 h. Quantification of (**E**) total LD area per cell and (**F**) LD size in AML12 cells transfected with control (siNT) or *Hrs*-targeting (si*Hrs*) siRNAs subjected to a pulse treatment of OA followed by a chase period in growth media (WD) with or without ATGLi. **G.** Immunoblot analysis for ATGL protein levels from siNT and si*Hrs* knockdown AML12 cells. **H.** Quantification of ATGL protein against GAPDH from the results in (**S2G**). Data represents the mean ± SD from three independent trials, where each dot represents an independent trial. The statistical significance is indicated by asterisks using a two-way ANOVA with a (**A, B**) Sidak’s, (**C, D**) Uncorrected Fisher’s LSD, or (**E, F**) Tukey’s post-hoc test, \**P* < 0.05, \*\**P* < 0.01, \*\*\**P* < 0.001, \*\*\*\**P* < 0.0001, n.s., not significant. Abbreviations: ATGL – adipose triglyceride lipase.

**Figure S3.** **A**. Confocal micrographs representing endogenous HRS by immunofluorescence (green) in AML12 cells where LDs are stained with ORO (red). **B.** Confocal micrographs representing AML12 cells transfected with an HRS-overexpression plasmid (red) where LDs are stained with MDH. **C.** Graph depicting organelle localization of HRS protein from mouse livers along a sucrose gradient.^21^

**Figure S4.** Quantification of total (**A**) 4E-BP1, (**B**) S6, (**C**) ULK1, and (**D**) S6 kinase in *Hrs*-knockdown (si*Hrs*) and or control (siNT) AML12 cells treated with or without Torin1 from (Figure 4H). The band intensities of each were normalized to the band intensities of ACTIN, followed by normalization of all groups against untreated siNT. Data represent the mean ± SD from at least two independent trials, where each dot represents an independent trial. The statistical significance is indicated by asterisks using a two-way ANOVA with an Uncorrected Fisher’s LSD post-hoc test, \*\*\*\**P* < 0.0001, n.s, not significant.

