## Supplemental Figure 1 for "The ESCRT-0 protein HRS regulates hepatocellular lipid droplet catabolism"

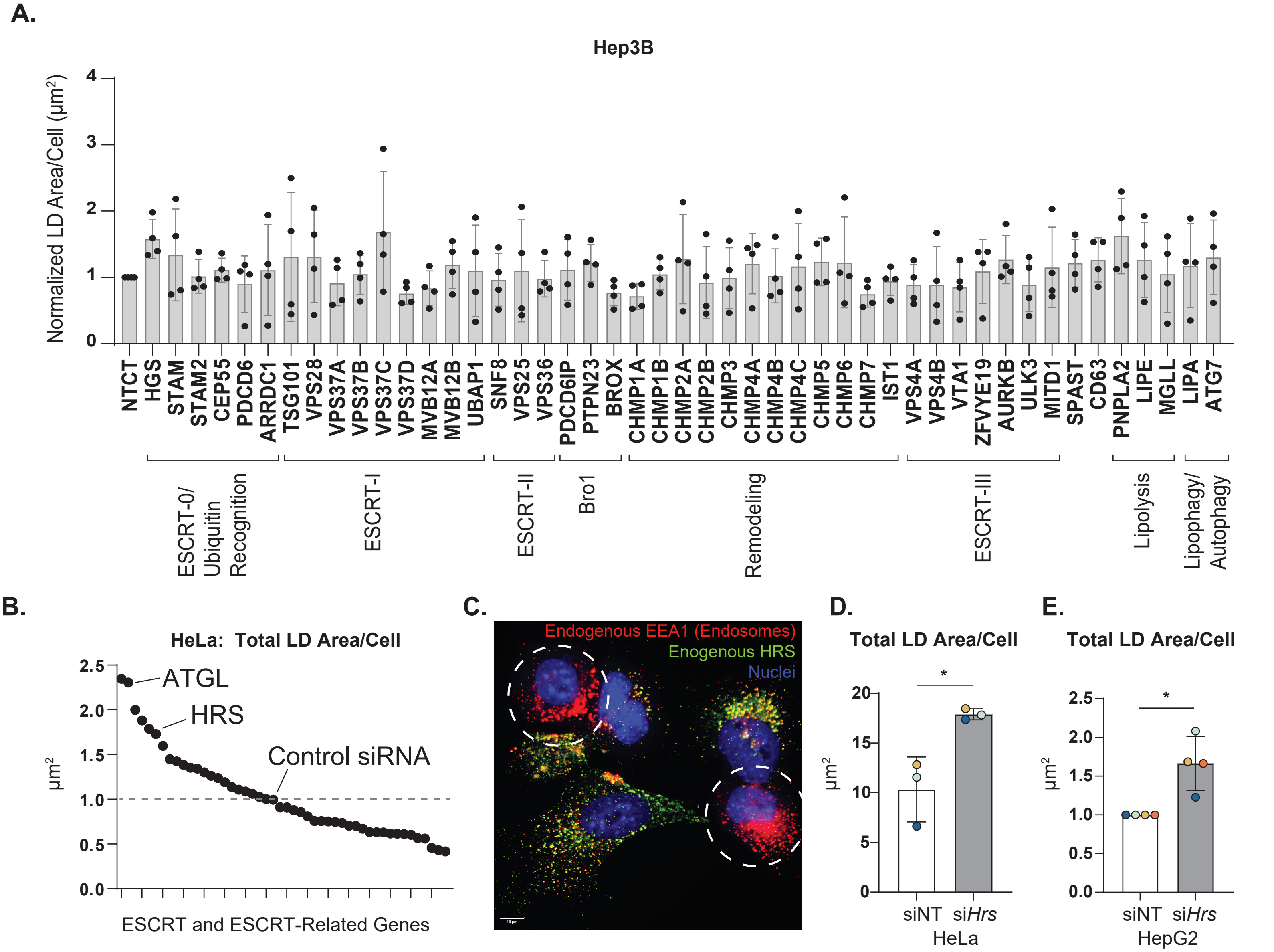


**Figure S1.**

A. Graph representing fold changes in total LD area per cell in Hep3B cells post knockdown of indicated ESCRT genes. B. Graph showing changes in total LD area per cell from RNAi screen of ESCRT and ESCRT-related genes in HeLa cells. C. Confocal micrographs of AML12 cells transfected with Hrs-targeting siRNA for 48 hours. The cells were then analyzed by immunofluorescence for early endosomes (EEA1, red), HRS (green), and DAPI-stained nuclei (blue). Quantification of total LD area per cell in (D) HeLa and (E) HepG2 cells treated with control (siNT) versus Hrs targeting (siHrs) siRNAs for 48 hours. Data represents the mean ± SD from at least three independent trials, where each dot represents an independent trial. The statistical significance is indicated by asterisks using a two-tailed Student's T-test for ratio-pairwise comparisons. *P < 0.05, **P < 0.01, n.s., not significant. Abbreviations: ESCRT – Endosomal Sorting Complex Required for Transport, LD: Lipid Droplet.
