## Supplemental Figure 2 for "The ESCRT-0 protein HRS regulates hepatocellular lipid droplet catabolism"

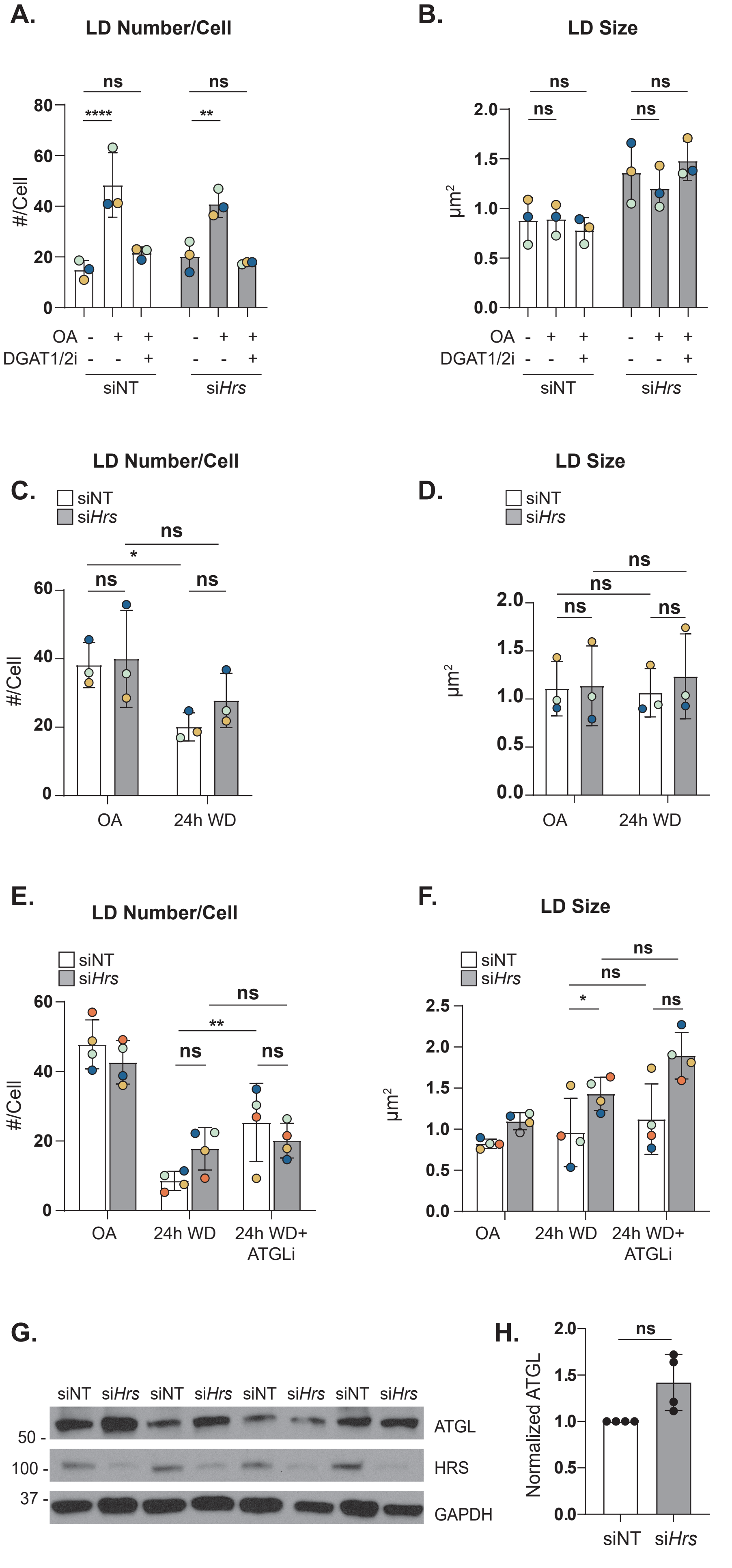


**Figure S2.**
Quantification of (A) LD number per cell and (B) LD size in AML12 cells transfected with control (siNT) or Hrs-targeting (siHrs) siRNAs and treated with OA, with or without DGAT1/2i. Quantification of (C) LD number per cell and (D) LD size in AML12 cells transfected with control (siNT) or Hrs-targeting (siHrs) siRNAs subjected to a pulse treatment of OA followed by a chase period in growth media without OA for 24 h. Quantification of (E) total LD area per cell and (F) LD size in AML12 cells transfected with control (siNT) or Hrs-targeting (siHrs) siRNAs subjected to a pulse treatment of OA followed by a chase period in growth media (WD) with or without ATGLi. G. Immunoblot analysis for ATGL protein levels from siNT and siHrs knockdown AML12 cells. H. Quantification of ATGL protein against GAPDH from the results in (S2G). Data represents the mean ± SD from three independent trials, where each dot represents an independent trial. The statistical significance is indicated by asterisks using a two-way ANOVA with a (A, B) Sidak's, (C, D) Uncorrected Fisher's LSD, or (E, F) Tukey's post-hoc test, *P < 0.05, **P < 0.01, ***P < 0.001, ****P < 0.0001, n.s., not significant. Abbreviations: ATGL – adipose triglyceride
