## Supplemental Figure 3 for "The ESCRT-0 protein HRS regulates hepatocellular lipid droplet catabolism"

A.

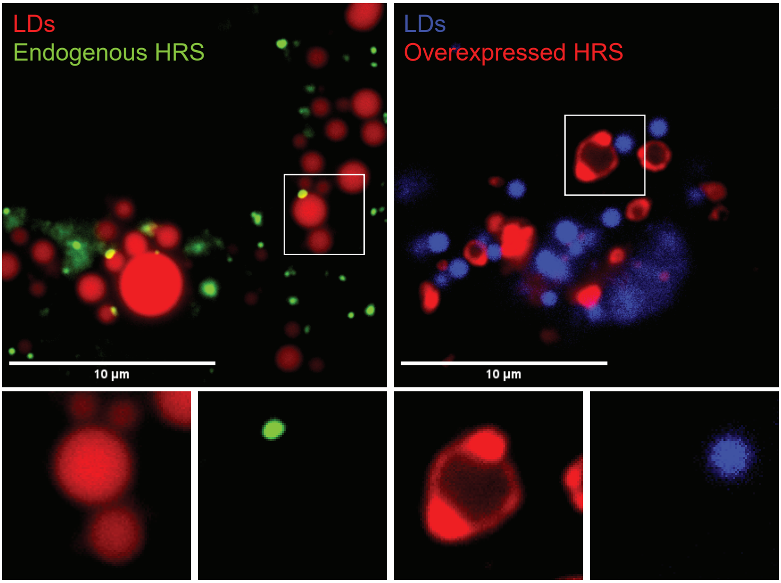


B.

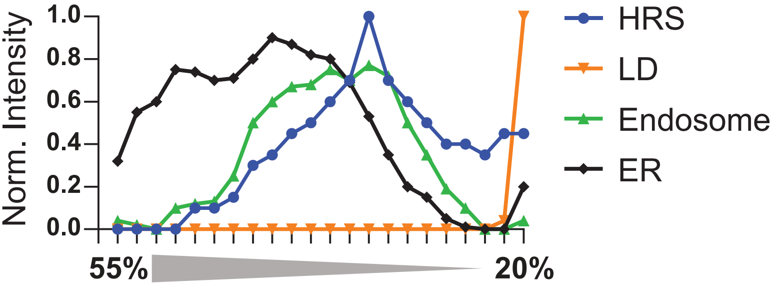


**Figure S3.**

A. Confocal micrographs representing endogenous HRS by immunofluorescence (green) in AML12 cells where LDs are stained with ORO (red). B. Confocal micrographs representing AML12 cells transfected with an HRS-overexpression plasmid (red) where LDs are stained with MDH. C. Graph depicting organelle localization of HRS protein from mouse livers along a sucrose gradient.[21]
