## Supplemental Figure 4 for "The ESCRT-0 protein HRS regulates hepatocellular lipid droplet catabolism"

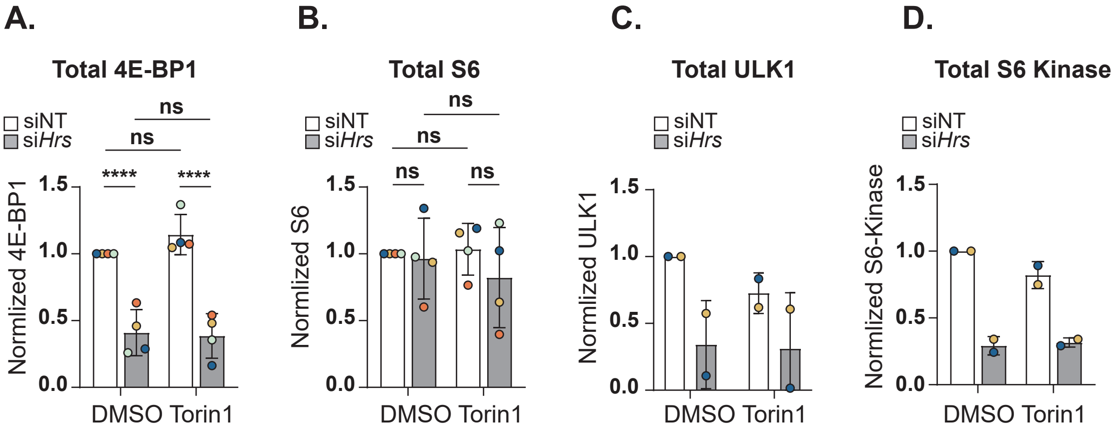


**Figure S4.**
Quantification of total (A) 4E-BP1, (B) S6, (C) ULK1, and (D) S6 kinase in Hrs-knockdown (siHrs) and or control (siNT) AML12 cells treated with or without Torin1 from (Figure 4H). The band intensities of each were normalized to the band intensities of ACTIN, followed by normalization of all groups against untreated siNT. Data represent the mean ± SD from at least two independent trials, where each dot represents an independent trial. The statistical significance is indicated by asterisks using a two-way ANOVA with an Uncorrected Fisher's LSD post-hoc test, ****$P < 0.0001$, n.s, not significant.
